# A precision medicine approach uncovers a unique signature of neutrophils in patients with brushite kidney stones

**DOI:** 10.1101/844142

**Authors:** Mohammad Shahidul Makki, Seth Winfree, James E. Lingeman, Frank Witzmann, Elaine M. Worcester, Amy Krambeck, Fred Coe, Andrew P. Evan, Sharon Bledsoe, Kristin Bergsland, Suraj Khochare, Daria Barwinska, James C. Williams, Tarek M. El-Achkar

## Abstract

**Background:** We have previously found that papillary histopathology differs greatly between calcium oxalate and brushite stone formers (SF); the latter have much more papillary mineral deposition, tubular cell injury and tissue fibrosis.

**Methods:** In this study, we applied unbiased orthogonal “omics” approaches on biopsied renal papillae and extracted stones from patients with brushite or calcium oxalate (CaOx) stones. Our goal was to discover stone type-specific molecular signatures to advance our understanding of the underlying pathogenesis.

**Results:** Brushite SF did not differ from CaOx SF with respect to metabolic risk factors for stones, but did exhibit increased tubule plugging in their papillae. Brushite SF had upregulation of inflammatory pathways in papillary tissue, and increased neutrophil markers in stone matrix compared to those with CaOx stones. Large-scale 3D tissue cytometry on renal papillary biopsies showed an increase in the number and density of neutrophils in the papillae of brushite vs. CaOx patients, thereby linking the observed inflammatory signatures to the neutrophils in the tissue. To explain how neutrophil proteins appear in the stone matrix, we measured neutrophil extracellular trap (NET) formation, NETosis, and found it significantly increased in the papillae of brushite compared to CaOx patients.

**Conclusions:** We show that increased neutrophil infiltration and NETosis is an unrecognized factor that differentiates brushite and CaOx SF, and may explain the markedly increased scarring and inflammation seen in the papillae of brushite patients. Given the increasing prevalence of brushite stones, the role of neutrophil activation in brushite stone formation requires further study.

## Introduction

Nephrolithiasis is a clinical syndrome that likely arises from multiple diseases each with a distinct pathobiology ^1^. For example, the papillary pathologies of patients who form idiopathic calcium oxalate (CaOx) stones differs from that of those whose stones contain brushite ^2^. The former show mainly interstitial plaque with very modest tubule plugging and no apparent tissue injury, whereas in the latter plugs are large and involve obvious tissue injury such as interstitial fibrosis and loss of tubule epithelial cells ^2^.

To date, the tissues from these two highly contrasting forms of kidney stone disease have not been studied using modern techniques to disclose mechanisms of injury. A role for inflammation and oxidative injury has been proposed based on animal models of crystal induced injury or *in vitro* studies ^3, 4^. However, definitive evidence for such a role in human disease is limited. A recent study by Taguchi et al, using transcriptomic studies on papillary tissue established an association between pro-inflammatory genes and Randall’s plaque development in CaOx stone formers (SF) ^5^. However, the papillary tissues of brushite SF, which have much more evidence for injury, have not been studied in similar fashion.

Significant advancement in molecular interrogation of tissue specimens presents a unique opportunity to uncover important pathways that are differentially activated in these two forms of kidney stone disease. These techniques such as transcriptomics, proteomics and large-scale quantitative 3-dimensional imaging can provide an unbiased and comprehensive approach towards discovery ^6–10^. We present here a detailed study contrasting CaOx and brushite clinical stone phenotypes using these orthogonal omics approaches.

Because these approaches are costly and require specific expertise of interpretation and integration our cohort is necessarily limited. Even so, our findings demonstrate that compared to CaOx SF, brushite SF have a unique signature of inflammation and neutrophil activation, which translates into increased abundance of neutrophils in the tissue and neutrophil proteins in the stones.

These findings were further extended by determining that these papillae are characterized by increased neutrophil extracellular trap formation (a process known as NETosis) ^11, 12^, whereby neutrophils expel their intracellular content. In addition to the pathophysiological implications of this unique neutrophil infiltration and activation in brushite stone formers, this work supports that NETosis, an important pathway hitherto undescribed in stone disease, could contribute to tissue injury in patients who form brushite stones.

## Results

### Patient characteristics

The two patient groups (Table 1) were similar in age (45.7 ± 14.2 vs 49.4 ± 12.7, mean ± SD, brushite and CaOx respectively, P=0.55), and sex distributions (89% vs 73% male, brushite and CaOx respectively p=0.59), but CaOx patients had higher BMI (25.7 ± 4.8 vs 32.6 ±7.4, brushite and CaOx, respectively, p<0.05). ESWL rates were similar (55.6% vs 45.5%, brushite and CaOx respectively, p=0.99). Few patients had comorbidities, and only one brushite patient had an infected urine culture. Stone cultures were negative (not shown). With respect to metabolic evaluation, excretions of calcium, oxalate and citrate, and supersaturation (SS) with respect to CaOx and CaP were also the same in both groups of patients (Table 2). This differs slightly from our prior work, in which calcium excretion and CaP SS were higher in brushite than CaOx patients ^13^.

**TABLE 1.** CLINICAL CHARACTERISTICS OF THE PATIENTS AND THE ASSAYS PERFORMED ON EACH SAMPLE. Mineral measurements were performed by volume through segmentation of 3D micro CT image stacks. CaOx, calcium oxalate; Ap, apatite; Br, brushite; ESWL, Extracorporel Shock Wave Lithotripsy; PN, Percutaneous Nephrolithotomy; URS, Uretroscopy; HTN, Hypertension; DP, Depression; DM, Diabetes mellitus; TD, Thyroid disease; CA, Cancer; GS, Gastric surgery; 3DTC, 3 dimensional tissue cytometry; Trans, transcriptomics; Prot, proteomics. * included 4.3% apatite, 2.8% octacalcium phosphate and 0.5% whitlockite. #, the type of surgery was unspecified.

| Pt | Stone Mineral (%) |  |  | Sex | Age at surgery (years) | Prior ESWL | Surgery type | Urine culture growth | BMI | Co-morbidities | Assays used on each tissue or stone specimen |  |  |  |
| --- | --- | --- | --- | --- | --- | --- | --- | --- | --- | --- | --- | --- | --- | --- |
|  | CaOx | Ap | Br |  |  |  |  |  |  |  | Tissue 3DTC | Tissue Trans | Stone Prot | Stone microCT |
| BRUSHITE PATIENTS |  |  |  |  |  |  |  |  |  |  |  |  |  |  |
| 1 | 4 | 8* | 88 | M | 56 | Yes | PN | No | 21.4 | No | - | - | + | + |
| 2 | 5 | 6 | 89 | M | 46 | Yes | PN | Yes | 24.8 | No | - | - | + | + |
| 3 | 5 | 15 | 80 | M | 54 | Yes | PN | No | 31.9 | No | - | - | + | + |
| 4 | 26 | 1 | 73 | M | 40 | No | PN | No | 28.6 | No | - | - | + | + |
| 5 | - | 6 | 94 | M | 43 | Yes | PN | No | 25.1 | No | + | + | - | + |
| 6 | 1 | 5 | 94 | M | 25 | No | PN, URS | No | 25.0 | No | + | + | - | + |
| 7 | 3 | 2 | 95 | F | 44 | No | URS | No | 19.1 | No | + | - | - | + |
| 8 | - | - | 100 | M | 74 | No | PN | NA | 22.4 | No | + | + | - | + |
| 9 | - | - | 100 | M | 29 | Yes | PN | No | 33.4 | No | + | + | + | + |
| CALCIUM OXALATE PATIENTS |  |  |  |  |  |  |  |  |  |  |  |  |  |  |
| 10 | 91 | 9 | - | M | 58 | No | PN | NA | 28.9 | No | - | - | + | + |
| 11 | 85 | 15 | - | M | 65 | No | PN | No | 36.2 | HTN,TD, CA | - | - | + | + |
| 12 | 94 | 6 | - | M | 21 | No | PN, URS | No | 27.8 | No | - | - | + | + |
| 13 | 82 | 18 | - | M | 47 | No | PN | No | 28.2 | HTN | + | - | + | + |
| 14 | 80 | 21 | - | F | 56 | Yes | PN | No | 25.5 | DM, TD, DP | + | + | - | + |
| 15 | 97 | 3 | - | F | 66 | Yes | PN, URS | NA | 40.3 | GS# | + | - | - | + |
| 16 | 85 | 15 | - | F | 50 | No | PN | No | 27.1 | No | + | - | - | + |
| 17 | 85 | 15 | - | M | 50 | No | PN | NA | 40.7 | No | + | + | - | + |
| 18 | 97 | 3 | - | M | 38 | Yes | PN | No | 47.4 | No | + | + | - | + |
| 19 | 99 | 1 | - | M | 53 | Yes | PN | No | 25.8 | No | - | + | - | + |
| 20 | 92 | 8 | - | M | 39 | Yes | PN | No | 31.0 | No | - | - | + | + |

**TABLE 2.** URINE AND SERUM ANALYTES. SS, supersaturation; CaOx, calcium oxalate; CaP, calcium phosphate as brushite.

| Patient | Urine volume (L/day) | Urine pH | Urine calcium (mg/day) | Urine citrate (mg/day) | Urine Oxalate | SS CaOx | SS CaP | Serum creatinine (mg/dL) | Serum calcium (mg/dL) |
| --- | --- | --- | --- | --- | --- | --- | --- | --- | --- |
| **BRUSHITE PATIENTS** |  |  |  |  |  |  |  |  |  |
| 1 | 3.19 | 6.69 | 252.6 | 671.0 | 52.3 | 5.55 | 0.87 | 1.12 | 10.13 |
| 2 | 1.94 | 6.37 | 371.6 | 697.8 | 41.8 | 8.55 | 3.32 | 0.83 | 9.54 |
| 3 | 1.63 | 7.74 | 122.4 | 567.6 | 57.5 | 6.18 | 2.06 | 0.83 | 9.78 |
| 4 | 0.97 | 6.28 | 260.3 | 958.5 | 26.0 | 9.94 | 4.20 | 1.30 | 9.93 |
| 5 | 2.13 | 6.67 | 295.0 | 225.0 | 47.0 | 7.52 | 2.69 | - | - |
| 6 | 1.70 | 6.31 | 279.4 | 297.4 | 38.6 | 7.62 | 2.19 | 0.75 | 9.39 |
| 7 | 1.58 | 6.25 | 290.3 | 447.0 | 29.4 | 8.91 | 2.22 | 0.91 | 9.56 |
| 8 | - | - | - | - | - | - | - | - | - |
| 9 | 1.71 | 6.52 | 191.7 | 949.7 | 65.5 | 8.84 | 2.29 | 0.92 | 9.00 |
| MEAN ± SD | 1.84 ± 0.68 | 6.31 ± 0.20 | 241.7 ± 63.0 | 648.7 ± 267.4 | 45.2 ± 14.6 | 7.80 ± 1.57 | 2.35 ± 0.98 | 0.97 ± 0.20 | 9.63 ± 0.40 |
| **CALCIUM OXALATE PATIENTS** |  |  |  |  |  |  |  |  |  |
| 10 | 1.69 | 5.97 | 145.5 | 425.0 | 45.6 | 6.42 | 0.81 | 0.62 | 9.47 |
| 11 | 2.11 | 6.51 | 256.7 | 278.9 | 50.0 | 8.03 | 2.32 | 0.79 | 8.83 |
| 12 | - | - | - | - | - | - | - | - | - |
| 13 | 1.33 | 6.01 | 315.9 | 881.6 | 30.2 | 8.65 | 3.73 | 1.04 | 9.24 |
| 14 | 2.79 | 6.55 | 494.3 | 899.9 | 40.0 | 8.17 | 2.64 | 0.72 | 9.65 |
| 15 | 1.55 | 6.28 | 91.1 | 253.7 | 56.1 | 7.89 | 0.77 | 0.92 | 7.99 |
| 16 | 0.91 | 5.89 | 310.5 | 972.3 | 28.5 | 16.11 | 3.52 | - | - |
| 17 | 1.55 | 5.63 | 293.8 | 670.5 | 51.4 | 15.51 | 0.96 | 0.92 | 8.31 |
| 18 | 1.25 | 5.45 | 528.5 | 949.8 | 64.9 | 26.47 | 2.25 | 0.84 | 9.42 |
| 19 | 1.59 | 5.90 | 479.2 | 940.2 | 41.4 | 12.52 | 2.23 | - | - |
| 20 | - | - | - | - | - | - | - | - | - |
| MEAN ± SD | 1.67 ± 0.51 | 6.22 ± 0.42 | 328.7 ± 144.8 | 697.0 ± 282.9 | 45.0 ± 11.2 | 6.14 ± 6.14 | 2.25 ± 1.10 | 0.83 ± 0.13 | 9.05 ± 0.61 |
| p value | 0.55 | 0.59 | 0.15 | 0.74 | 0.97 | 0.11 | 0.84 | 0.15 | 0.07 |

### Kidney stone characterization

By micro – CT analysis, CaOx stones were heterogeneous. In a majority, mineral type was calcium oxalate monohydrate (COM), and all contained some apatite (Figure.1 top and Table 1). The average specimen composition (by volume) for the CaOx stone formers was 58.6± 30.1% COM, 31.1± 29.1% calcium oxalate dihydrate (COD), and 10.3± 6.4% apatite. In contrast, the brushite specimens were homogeneous (95.4± 4.7% brushite, 1.7± 3.6% COD, and 2.9± 4.1% apatite (Figure.1 bottom and Table 1).

**Fig. 1.**
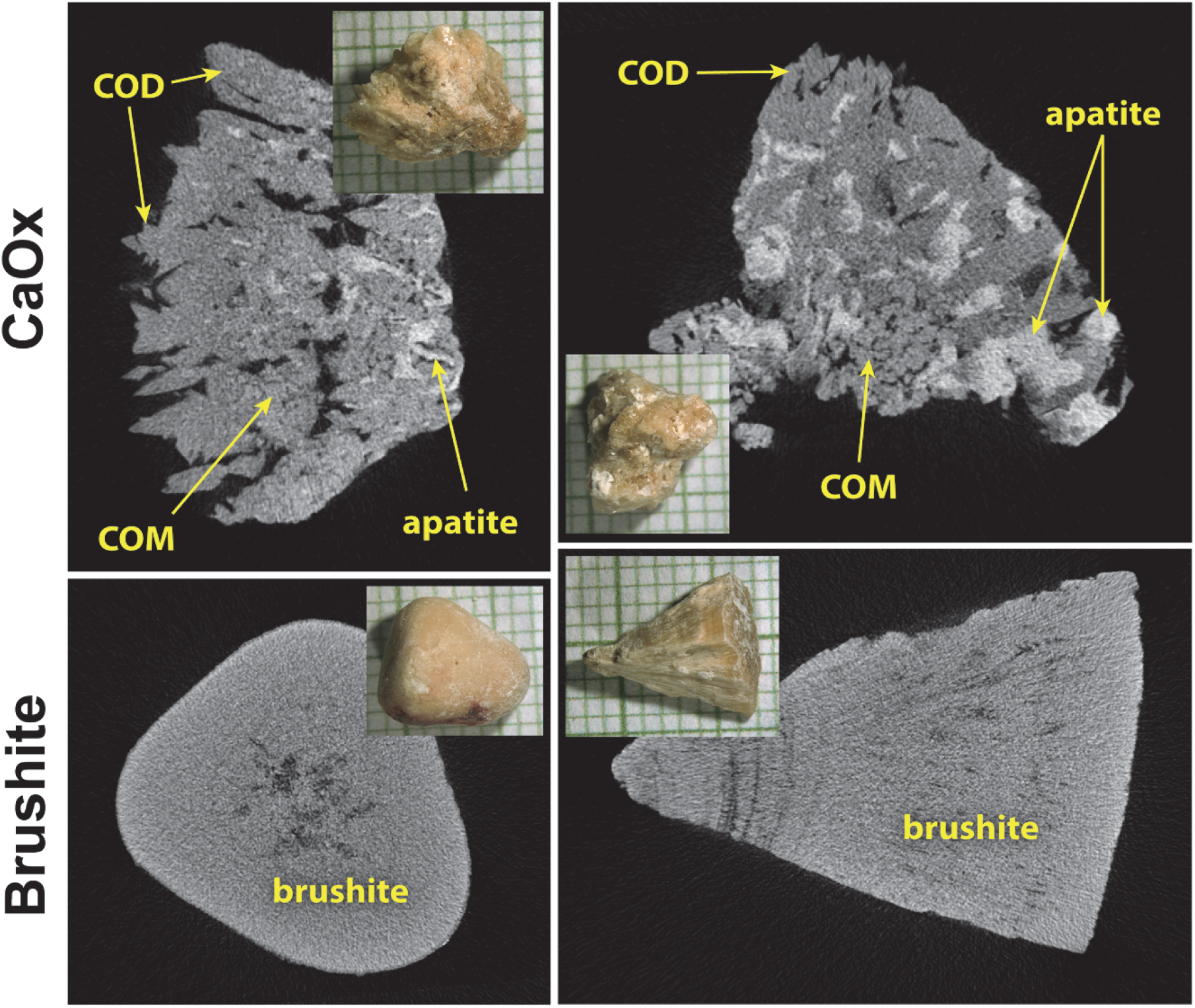
Micro CT of kidney stones. Kidney stones are shown in insets, on mm paper. Upper panels show two typical calcium oxalate (CaOx) specimens, both of which show polyhedral crystals of calcium oxalate dihydrate (COD) at the surface, calcium oxalate monohydrate (COM, which is slightly more X-ray dense than COD), and apatite, which is the most X-ray dense of the minerals commonly found in stones. Specimen in the upper right contained the largest fraction of apatite of any of the specimens used (21% apatite by volume). The lower panels show brushite specimens, both of which were quite pure, which was typical for all the brushite samples.

### Surgical and tissue phenotyping

Surgery disclosed variable amounts of plugging and Randall’s plaque in both groups of patients (Figure 2, panels a and b), which were confirmed histologically (Figure 2, panels c and d). Plugs and plaque in both types of stone formers had been shown to be composed of apatite ^13^. Papillary grading (Table 3) showed higher plugging but lower Randall’s plaque scores in brushite vs. CaOx (Fig. 2E and F), which is consistent with earlier reports ^13^. When assessed using surface area measurements (Table 3) only the amount of plugging differed between stone types.

**Fig. 2.**
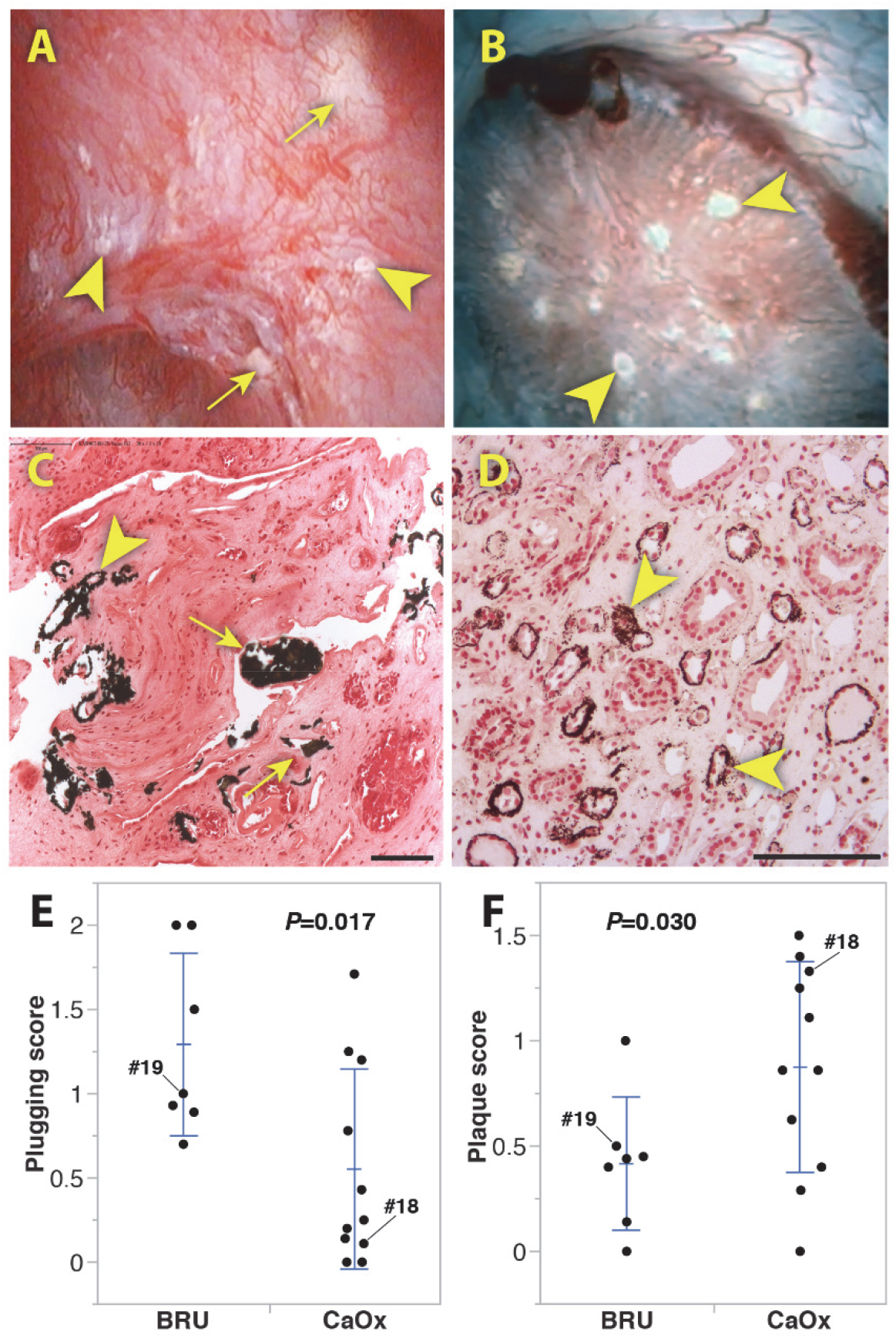
Endoscopic and histologic analysis of brushite and CaOx papillae. A and B show endoscopic appearance of papillae in brushite (patient #8) and CaOx (patient #20) patients, respectively. C and D show histological images (Yasue stain) of biopsies from these same patients. Arrowheads mark Randall’s plaque, and arrows point to regions of ductal plugging; bars in lower right indicate 100 μm. E and F show papillary scores for plugging and Randall’s plaque for all the patients except for 2 brushite patients, one of which had advanced scarring precluding scoring, and another with low quality endoscopic video. In the scatter plots presented, each point represents the mean score per category from a single patient. Error bars show standard deviation around the group mean. *P* values shown are from t-tests with Welch correction. Brushite (BRU).

**TABLE 3.** PAPILLARY GRADING SCORES. # papillae, number of papillae from each patient that were able to be graded. Plugging, Pitting, LoC (loss of papillary contour), and RP (Randall’s plaque) are mean scores for papillae graded ^43^. % plugging and % RP are obtained by measuring surface areas of plugging (yellow plaque) or RP on still frames from each papilla ^55^. P-values are t-tests assuming unequal variances, comparing brushite and CaOx

| Patient | # papillae | Plugging | Pitting | LoC | RP | % plugging | % RP |
| --- | --- | --- | --- | --- | --- | --- | --- |
| **BRUSHITE PATIENTS** |  |  |  |  |  |  |  |
| 1 | 2 | 1.5 | 0 | 0.5 | 1 | 2.39 | 2.70 |
| 2 | 9 | 0.89 | 0.78 | 0.56 | 0.44 | 0.97 | 3.26 |
| 3 | 15 | 0.93 | 0.47 | 1.47 | 0.4 | 0.71 | 0.64 |
| 4 | all papillae scarred with no recognizable features |  |  |  |  |  |  |
| 5 | 6 | 2 | 0.67 | 0.67 | 0 | 3.24 | 0.04 |
| 6 | 20 | 0.7 | 0.15 | 0.4 | 0.45 | 0.02 | 2.40 |
| 7 | 7 | 2 | 1.71 | 0.57 | 0.14 | 3.57 | 0.50 |
| 8 | 6 | 1 | 0.17 | 0.17 | 0.5 | 1.52 | 0.50 |
| 9 | video too dark to discern details for grading |  |  |  |  |  |  |
| **MEAN ± SD** | 9.3±6.1 | 1.3±0.5 | 0.6±0.6 | 0.6±0.4 | 0.4±0.3 | 1.8±1.3 | 1.4±1.3 |
| **CALCIUM OXALATE PATIENTS** |  |  |  |  |  |  |  |
| 10 | 6 | 0 | 0 | 0 | 1.5 | 0 | 4.51 |
| 11 | 4 | 0.25 | 0.5 | 0.25 | 1.25 | 0.06 | 1.47 |
| 12 | 8 | 0 | 0.375 | 0 | 0.625 | 0 | 4.39 |
| 13 | 7 | 0.14 | 0 | 0.29 | 0.29 | 0.05 | 0.45 |
| 14 | 5 | 1.2 | 0 | 0 | 0.4 | 0.79 | 1.29 |
| 15 | 9 | 0.78 | 0.33 | 0 | 1.11 | 0 | 2.68 |
| 16 | 7 | 0.43 | 0.43 | 0.71 | 0.86 | 0.15 | 2.17 |
| 17 | 7 | 1.71 | 0.43 | 0.14 | 0.86 | 3.20 | 1.77 |
| 18 | 5 | 0.2 | 0.4 | 0.8 | 1.4 | 0.052 | 3.34 |
| 19 | 4 | 1.25 | 0.25 | 0 | 0 | 0.47 | 0.15 |
| 20 | 9 | 0.11 | 0 | 0.33 | 1.33 | 0 | 6.56 |
| **MEAN ± SD** | 6.5±1.8 | 0.6±0.6 | 0.3±0.2 | 0.2±0.3 | 0.9±0.5 | 0.4±1.0 | 2.6±1.9 |
| P-value | 0.277 | **0.017** | 0.207 | 0.052 | **0.030** | **0.044** | 0.141 |

### Transcriptomic profiling uncovers an immune activation molecular response in brushite papillary biopsies

We extracted RNA from cryopreserved papillae (n=4 each group), and performed RNA sequencing analysis as described in the methods. Differential expression analysis is presented in supplemental data file 1. Bioinformatic pathway analysis on differentially expressed genes showed that the pathway with most significant gene clustering (p<0.01, Fisher’s exact test) and highest z-score for activation (z score >2) was that of pathogen recognition by the immune system (Fig. 3A-C), which included pro-inflammatory genes such as TLR4, TLR8, IL17D and PIK3R1. These were among the top upregulated genes in the brushite vs. CaOx papillae (Fig. 3C). Additional disease pathway and network analysis also confirmed a heightened injury and inflammatory activation signature in brushite vs. CaOx (Fig. 3D and 3E). Notably, many genes in the inflammatory pathways derived from this analysis are involved in inflammation and neutrophil activation (Fig. 3D and 3E).

**Fig. 3.**
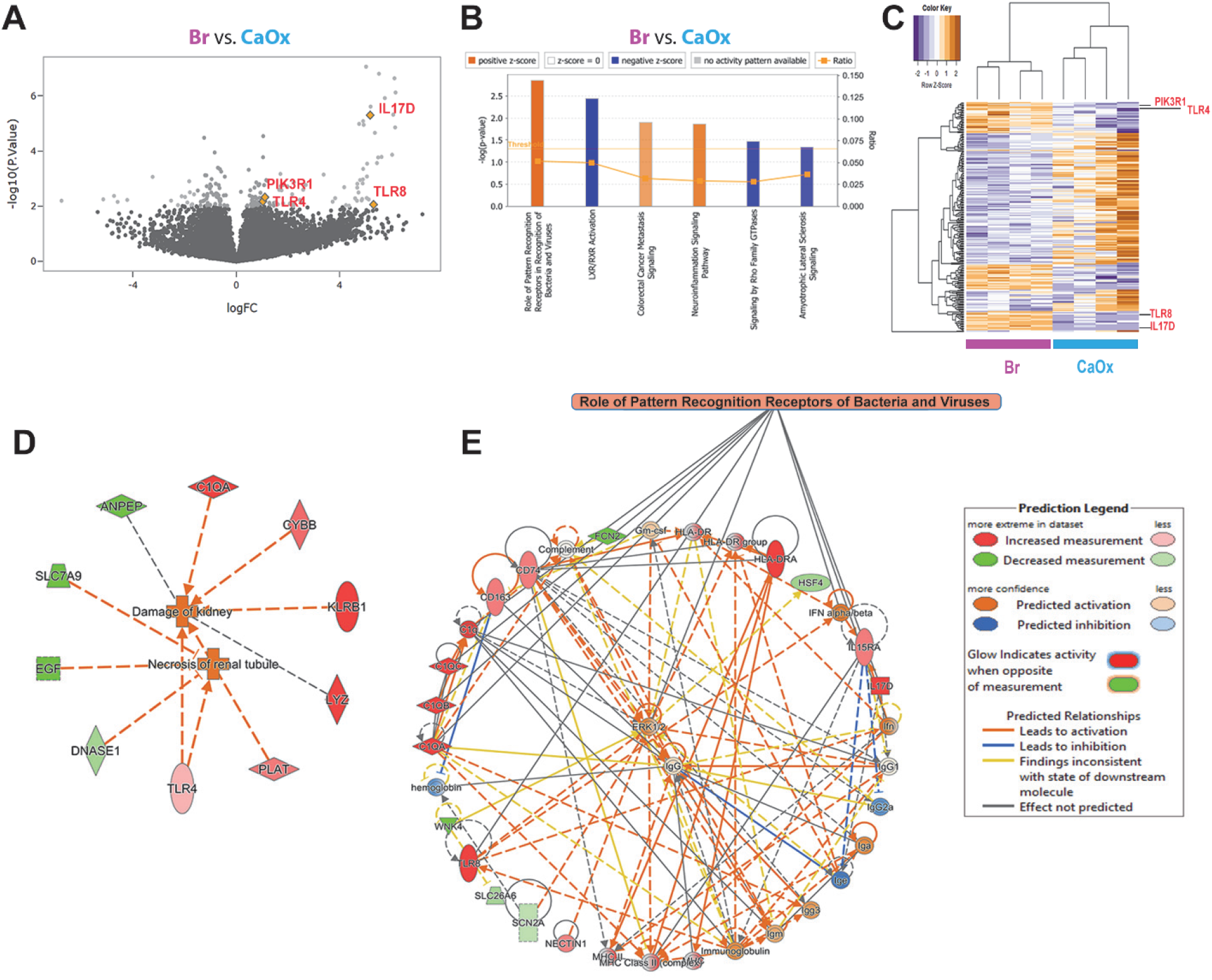
Transcriptomic analysis of papillary biopsies. (A) Total RNA obtained from four CaOx and four brushite papillary biopsies were individually sequenced. Volcano plot was made using negative log10 transformed p-values against the log ratio using values from brushite vs CaOx. Few genes that are part of the top significantly upregulated pathway are shown. (B) Differentially expressed genes as described in Methods (brushite vs CaOx) were analyzed for pathway enrichment using IPA, and significantly altered pathways (p<0.05) are shown with absolute z score >1 (activation: orange; inhibition: blue). The top pathway was: “Role of pathogen recognition receptors in recognition of bacteria and viruses” with p=0.00142 and z score for activation of 2.236. (C) Unsupervised hierarchical clustering was constructed for the top 300 genes sorted by p value, showing also genes from the top upregulated pathway. (D) Disease pathway analysis differentially activated in brushite vs. CaOx, supporting heightened injury in the brushite papillae. (E) Signaling pathway analysis consistent with inflammatory activation signature in brushite vs. CaOx. Color code is indicated in the legend. The orange colored boxes indicate predicted activation, based on also on IPA.

### Label-free quantitative mass spectrometry (LFQMS) of kidney stones reveals a unique signature of neutrophil activation in brushite stones

In stones from 5 patients from each group, we identified 1947 proteins available across the sample types (supplemental file S2). Bio-informatics analysis on differentially abundant proteins in brushite vs. CaOx identified neutrophil-specific pathways such as leukocyte extravasation and IL-18 signaling among the top significantly enriched pathways (p< 0.01, Fisher’s exact test) with highest activation z-scores (>2) (Figure. 4). We identified 96 proteins in our data as definite neutrophil proteins (Supplemental data file 4) ^14–20^. Heat map visualization (Fig. 4C) indicated clustering and increased abundance of neutrophil activation proteins [e.g. Myeloperoxidase (MPO) fold change 5.53, FDR adjusted p value = 0.006), vascular non-inflammatory molecule-2 (VNN-2), fold change 3.52, FDR adjusted p value = 0.02)] in brushite compared to CaOx stone.

**Fig 4.**
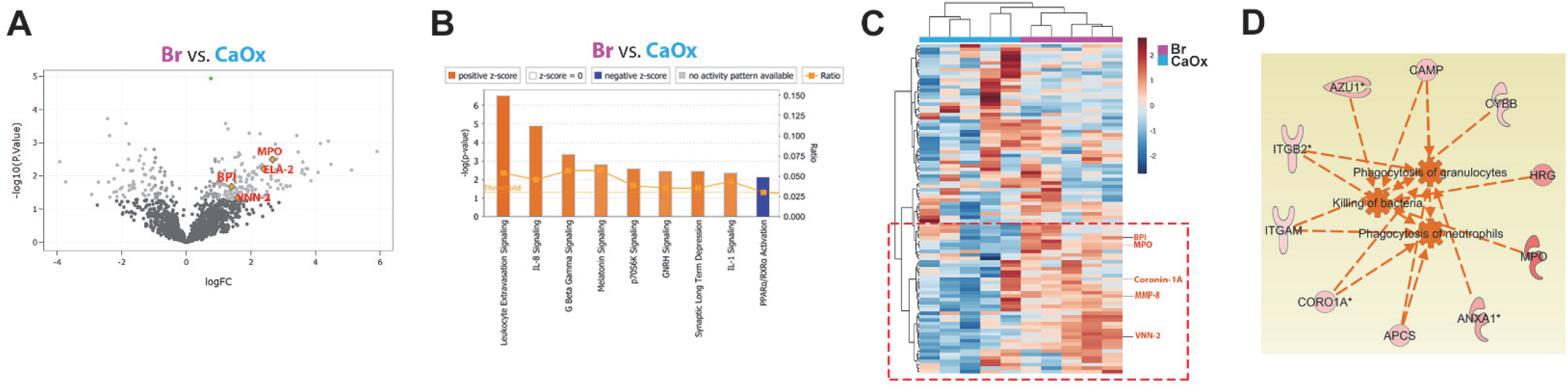
Proteomic analysis of kidney stones. (A) Volcano plot showing p values (−log10) versus protein ratio of brushite vs CaOx for all 1716 proteins fulfilling strict quantification criteria. Gene name of few proteins involved in inflammatory signaling are indicated with MPO and VNN-2 significantly increased by FDR. (B) The top canonical pathways enriched by IPA analysis are shown in a design similar to Fig 3. (C) Unsupervised hierarchical clustering of neutrophil specific proteins is shown, and the red box underscores proteins involved in neutrophil activity, and few examples are indicated to the right. (D) disease pathways predicted by IPA. The proteins indicated are related to activation of neutrophil specific pathways such as “phagocytosis of granulocytes”, “killing of bacteria”, “phagocytosis of neutrophils” in brushite vs. CaOx. The same color codes from the legend used in Fig 3. apply here.

### Large scale 3D tissue cytometry maps the molecular signature of neutrophil activation to cell biology *in situ*

Large scale 3D confocal imaging was performed on frozen papillary biopsies (brushite and CaOx patients), stained for MPO, CD68, AQP1 and DAPI, to label neutrophils ^21^, activated macrophages ^22^, thin descending limbs ^23^, and nuclei, respectively. Quantitative 3D tissue cytometry was performed using the volumetric tissue cytometry and analysis software (VTEA) (Figure 5A and B) ^7^.

**Fig. 5.**
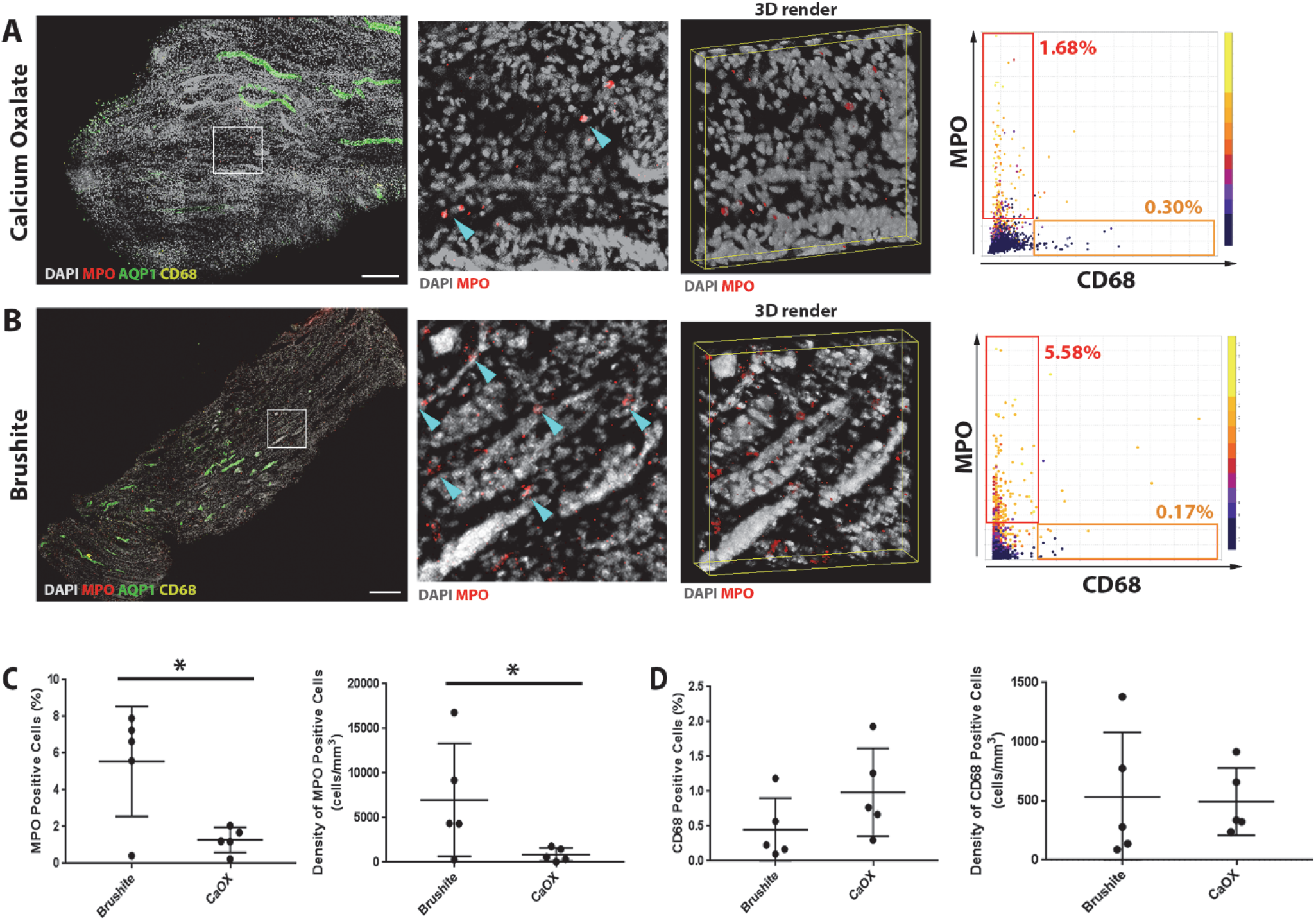
Large scale 3D imaging and tissue cytometry analysis of inflammatory cells in papillary biopsies. Fifty micron thick CaOx (A) or Brushite (B) papilla biopsy sections were immuno-labeled with anti-MPO (red), anti-AQP1 (green) and anti-CD68 (yellow) antibodies and DAPI (gray). At left, large-scale maximum projection images of entire biopsies with insets as indicated and a 3D rendering. Arrowheads indicate neutrophils. At right, representative scatter plots from VTEA are shown with the percentage of positive cells in the gated areas as indicated. Scale bar = 100 μm. Quantifications obtained from VTEA analysis were plotted (C and D) showing percent (out of total number of cells) and density of MPO+ and CD68+ cells, respectively. Each dot represents the value from a single patient’s biopsy. *Two-tailed t-test, p<0.05.

Both neutrophil abundance (5.55 ± 2.68% vs. 1.27 ± 0.61%, p<0.05) and density (6.99×10^3^ ± 5.6×10^3^/mm^3^ vs. 0.87×10^3^ ±0.67×10^3^/mm^3^; p<0.05) were significantly higher in brushite vs. CaOx biopsies, respectively (Figure. 5C). Importantly, we did not notice any difference in the abundance or density of CD68+ cells in brushite compared to CaOx (Figure. 5D). Therefore, these findings confirm a specific increase in neutrophil cell infiltration in the papillae of brushite patients.

Since plugging could cause inflammation, we asked if there is a link between neutrophil infiltration and the extent of papillary plugging. Figure 6A shows no correlation between neutrophil infiltration and plugging or Randall’s plaque formation. Furthermore, spatial correlation analysis, which was performed in one of the brushite biopsies, showed no distance correlation of neutrophil distribution to the location of the collecting duct (Figure 6B, C). These findings support that increased neutrophil infiltration in the brushite papillae is diffuse and does not cluster around a particular nidus of pathology.

**Fig. 6.**
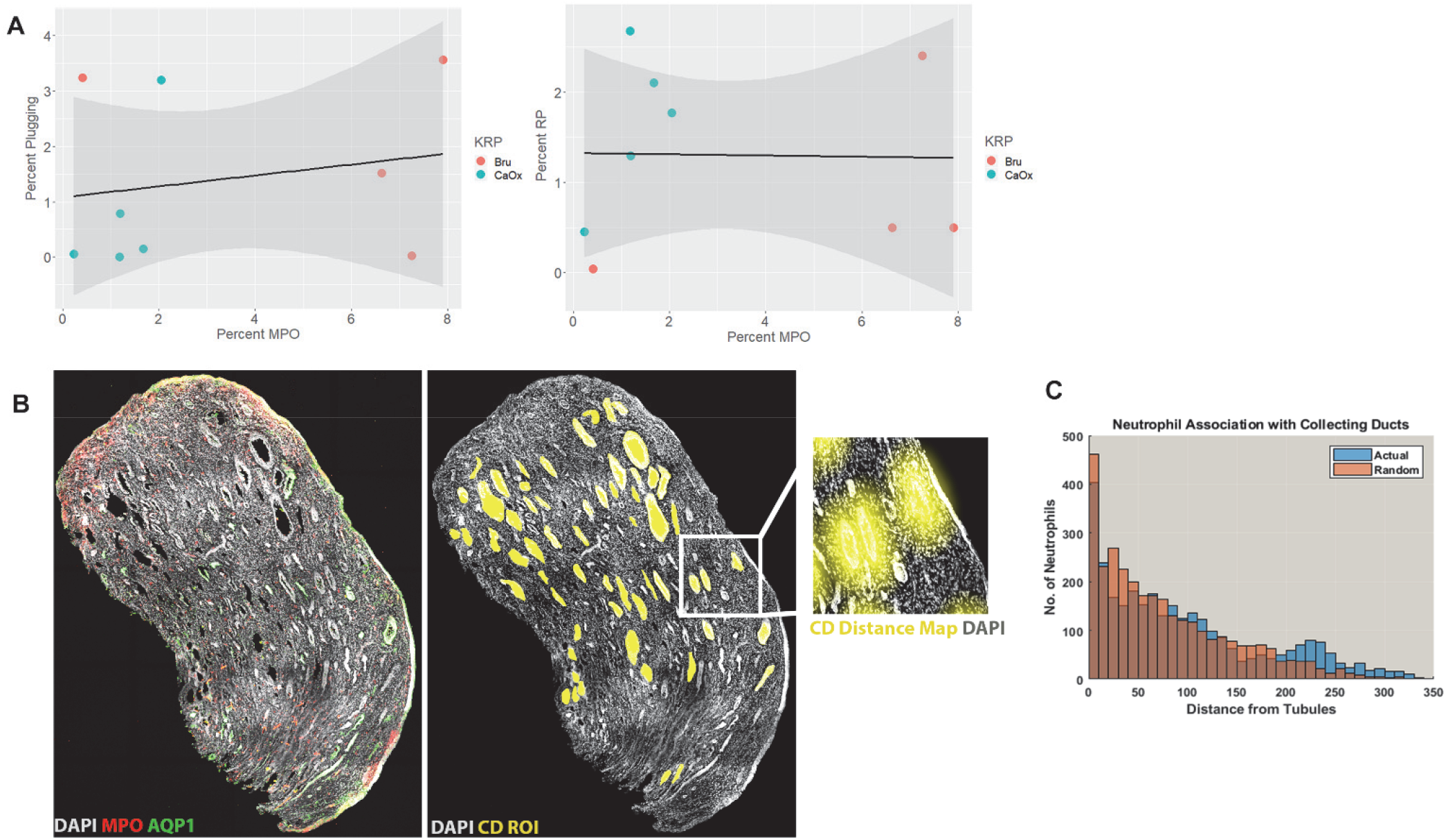
Correlation analysis between neutrophil distribution and papillary pathology. (A) Linear regression analysis was performed between neutrophil abundance in Brushite (Bru) and CaOx papilla biopsies compared to papillary endoscopic scoring of plugging (left) or Randall’s Plaque (RP) (right). No correlation was observed between these events (*p*=0.60 and 0.94, respectively). (B) Maximum projection of a papillary biopsy labeled with MPO (neutrophils) or AQP1 (thin descending limb) shows diffuse pattern of neutrophil distribution (left). Region of interests (ROI) marking collecting ducts were manually drawn (yellow) and distance maps were built for every ROI (example of a distance map shown in the enlarged image in inset). The positions of all the neutrophils surveyed using VTEA tissue cytometry analysis were correlated to locations on the distance maps. Neutrophil distance distribution was nearly identical to a randomly generated distribution (Gaussian), shown in the histograms depicted in (C) (p=1.0), suggesting no spatial correlation between neutrophil distribution and collecting ducts.

### Increased NETosis in the papilla could explain the abundance of neutrophil proteins in brushite stones

In the brushite tissue specimens many nuclei of neutrophils were poorly demarcated, which led us to hypothesize the presence of NETosis. Because citrullination of histone is a marker of NETosis (Figure 7A), we probed papillary biopsies for citrulline and performed 3D imaging and cytometry to quantify NETosis (Figure 7B-C Left). Using VTEA analysis, we confirmed that NETosis (citrulline+ neutrophils) was significantly increased in brushite vs. CaOx papillae (0.18 ± 0.05% vs. 0.05 ± 0.02%; p<0.05) (Figure 7D). Interestingly, there was also a significant increase in the citrullination of cells other than neutrophils in the brushite papillae (Figure 7E), which is also consistent with an inflammatory state ^24^.

**Fig. 7.**
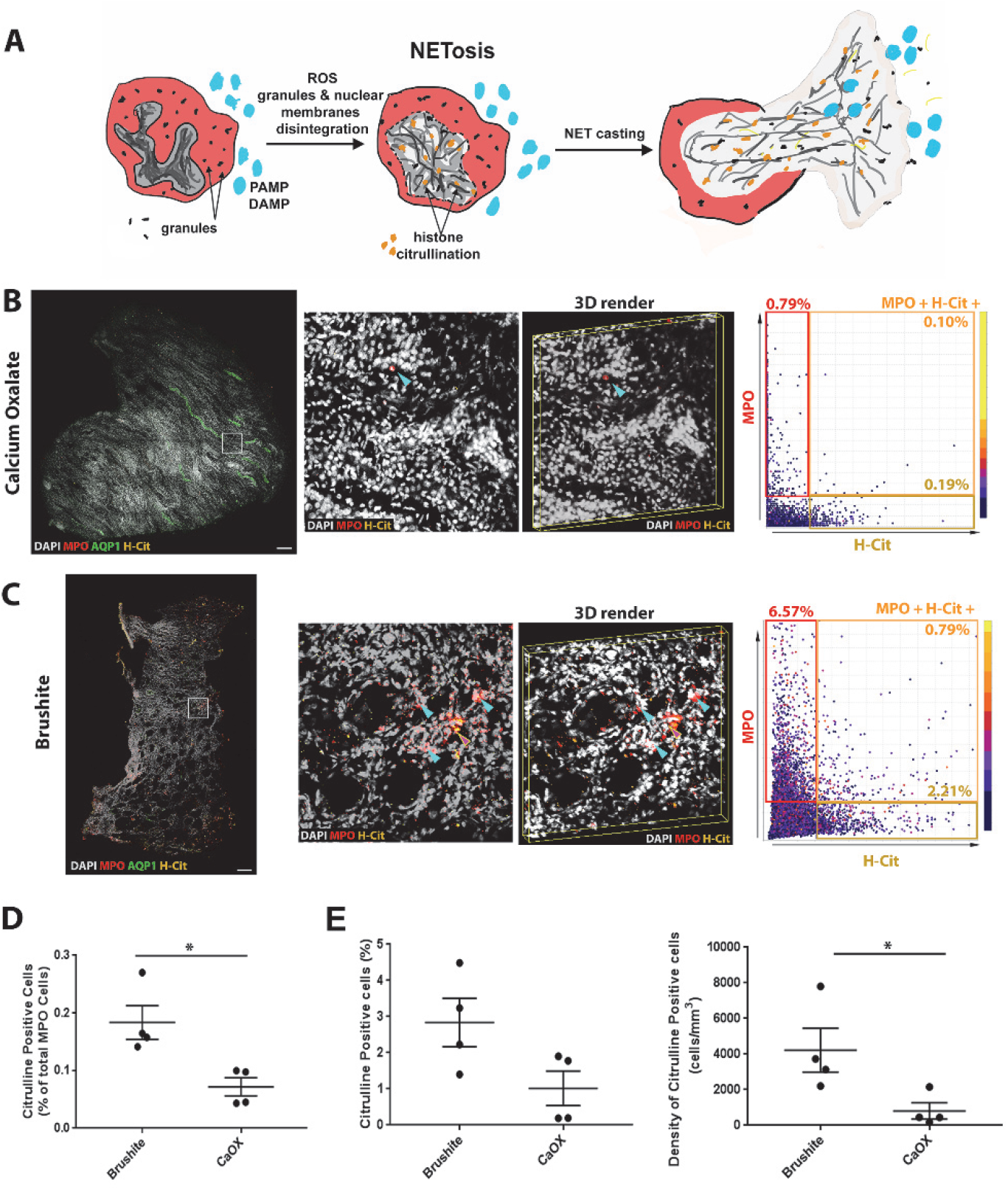
Neutrophil Extracellular Trap formation (NETosis) in papillary biopsies. (A) Figure shows an illustration of neutrophils undergoing NETosis, triggered by invading pathogen associated molecular patterns (PAMP) or disease associated molecular patterns (DAMP). NETosis is initiated by intracellular calcium influx and reactive oxygen species (ROS) production. The intracellular signaling induces histone citrullination (yellow), chromatin condensation and disintegration of nuclear and granular membranes which lead to the formation of NETs. These NETs consist of DNA packed with nuclear (e.g. histones) and granule (e.g. MPO, elastase) proteins. (B-C) Fifty micron thick CaOx (B) or Brushite (C) papilla biopsy sections were immuno-labeled with anti-MPO (red), anti-AQP1 (green) and anti-citrullinated histone (H-Cit) (yellow) antibodies and DAPI (gray). At left, large-scale maximum projection images of entire biopsies with insets as indicated and a 3D rendering. Blue arrowheads indicate neutrophils, whereas red arrowhead point to neutrophils that also stain for H-Cit. At right, representative scatter plots from VTEA are shown with the percentage of positive cells in the gated areas as indicated. Scale bar = 200 μm. (D-E) Quantifications obtained from VTEA analysis were plotted (D and E), showing the percentage of H-Cit+ neutrophils undergoing NETosis (D) and percentage and density of non-neutrophil cells with citrullinated histones. Each dot represents the value from a single patient’s biopsy. *Two-tailed t-test, p<0.05.

## Discussion

In this work, we determined that patients with different types of stone disease exhibit distinct cellular and molecular signatures. Our main finding is that tissue from brushite, but not CaOx stone formers, shows abundant neutrophil activation, including NETosis, a neutrophil response to bacteria and other potential pathogens. In addition, neutrophil derived proteins are abundant in brushite, but not CaOx, stone matrix.

The marked difference between brushite and CaOx stone formers with respect to papillary and stone matrix findings is especially striking given the fact that in this series demographic and metabolic characteristics of the two patient groups are so similar. Brushite stones carry a worse prognosis than CaOx stones in certain respects, as brushite stones have increased risk of recurrence ^25^, are frequently large and bilateral ^26^, and require more extra-corporeal lithotripsy procedures ^27^ compared to CaOx stones. Of concern, brushite stones appear to be increasing in prevalence ^28^ including among pediatric stone formers ^29^. In some cases, patients who initially make CaOx stones may later convert to making stones formed predominantly of calcium phosphate ^30, 31^. In a prior publication we documented that a significant fraction of our brushite SF had initially made stones composed mainly of CaOx ^13^. The association of papillary inflammatory activation with brushite stone formation suggests that conversion may involve more than just simple factors related to relative supersaturation with respect to calcium phosphate, as some have proposed ^32^. At this time it is not possible to tell whether the papillary inflammatory findings preceded brushite stone formation or followed it, however the finding of neutrophil related proteins preferentially in brushite stone matrix suggests the former.

The signal stimulating the inflammatory response in brushite SF is unknown. Crystals are known to induce inflammation, and cause NET formation when exposed to neutrophils *in vitro* ^3, 33, 34^. Although crystal-induced inflammation is quite possible in the brushite papilla, the widespread infiltration and the lack of clustering of neutrophils in specific areas of deposits argue against this, as does the fact that there was no significant relationship between tubule plugging and the signal for NETosis in the patients as a whole (Figure 6). Ascending infections can trigger neutrophil infiltration and NETosis ^11^. Brushite patients may have subclinical infections of papillary tissue, thereby linking infection to brushite stone formation. Although there were no obvious signs of increased urinary infection rate in the brushite group, not all bacteria residing in urine may be amenable to detection by typical culture methods ^35^. Additional studies are needed to test this hypothesis, knowing that the therapeutic implications of a positive finding would be immense: treating an infection to prevent stone formation.

Metabolic abnormalities such as oxidative stress deserve consideration ^36, 37^. Oxidative stress can trigger a pro-inflammatory signaling that promotes cell injury, apoptosis and inflammation. Our findings, especially of increased overall citrullination (Fig. 7E) and altered expression of multiple histone genes in brushite papillary tissue raise this possibility ^38^, and will require future testing. Likewise, injury from SWL could have led to our findings, but both groups had similar exposure to SWL, making this a less plausible explanation.

The released neutrophil proteins were a major component of the brushite stone matrix. Whether neutrophil proteins are a nidus of brushite crystal nucleation, or stabilize brushite crystals to prevent their spontaneous transformation to apatite ^13^ are hypotheses beyond the present study. At present it is not known if brushite stone formers are unique in having these inflammatory findings in papillary tissue. Other types of stones are associated with tubule plugs and scarring ^1^, and future studies will be needed to determine if similar inflammatory pathways are upregulated in these stone phenotypes as well.

From a methodological standpoint, this study demonstrates how to integrate clinical phenotyping with large multi-modal omics datasets and quantitative 3D imaging and tissue cytometry. Characterization of stone composition and tissue mineral deposits were confirmed with high precision using micro CT. The ability to map pathways discovered using an unbiased approach to the cell biology within the tissue, is a demonstration of the strength and potential uses of such an integrated multi-dimensional approach.

Our study has limitations. The representativeness of the small cohort of patients and the generalizability of our findings are weaknesses, pointing to the need to validate these findings in a larger cohort. However, the ranges of many of the clinical variables in the patients from our study, such as urine pH and mineral supersaturation, were comparable to other studies ^13, 26, 39^, suggesting that the cohort used here is comprised of patients with common stone presentations. The fact that not all interrogation techniques could be performed on specimens from the same patients is another limitation. However, the fact that the same findings were detected in different patients using orthogonal techniques could be viewed as a strength.

Despite the small number of patients, the depth provided by the molecular interrogation techniques was still sufficient to clearly detect specific molecular signals. The demonstration that this approach can extract meaningful results from a small cohort of patients is another strength of this study. Although we did not observe differences in all the proteins associated with neutrophils between the two types of stones, it is unlikely that all proteins released in the urine are equally incorporated into stones. Also, some neutrophil proteins like S-100 are made by other cells such as collecting duct cells, which will mask the specific contribution of neutrophils ^40^.

In conclusion, we demonstrate the presence of a stone-specific molecular signature of neutrophil activation and NETosis, that informs on the pathogenesis of brushite stone formation. In the process, we showcase the strength of a multi-modal approach consisting of thorough clinical phenotyping, integrated with high resolution cellular and molecular interrogation, and linked to the biology *in situ* using large scale 3D imaging. In addition to the overall implications for the pathogenesis of nephrolithiasis, our study could serve as a model and a proof of concept for large precision medicine initiatives that are seeking to implement such multidimensional approaches to better classify and understand human disease.

## Methods

### Subjects and Specimen Collection

Patients who were undergoing either percutaneous nephrolithotomy or ureteroscopic removal of renal stones were consented ^41, 42^. Patients were included in this study if they had CaOx or brushite stones, defined as stone mineral content >50% of the specified material, and also available specimens (stone or tissue) that could be used experimentally. The entire endoscopic procedure was recorded on video, papillary visual appearance was graded as previously described ^43^ and papillary biopsies were taken when possible. The study was approved by the Institutional Review Board Committee for Indiana University Health Partners (#98-073). Specific details of the specimens used for each patient are described in Table 1.

Stones were rinsed in normal saline and air dried at room temperature (RT) for further analysis by micro CT ^44^, using voxel sizes of 3-10 μm for the final reconstructions. Micro CT stone analysis was always confirmed using conventional Fourier transform infrared (FT-IR) spectroscopic method ^44^.

### Clinical Laboratory Studies

All subjects collected two 24-hour urine samples post-operatively while eating a free choice diet and off medications that could affect stone formation. Stone risk analytes were measured and urine supersaturation (SS) calculated using methods detailed elsewhere ^42, 45^. The mean of the two samples is reported in Table 2. Routine blood measurements were made for clinical purposes.

### Tissue

Whenever possible, papillary biopsies included both those that were immediately fixed in 4% paraformaldehyde (PFA, buffered to pH 7.4) or immediately immersed in OCT (optimal cutting temperature) medium and frozen on dry ice. Biopsies fixed in PFA were embedded in paraffin, sectioned and stained. Biopsies that were frozen were cut in a cryostat (Leica Biosystems, Wefzlar Germany) into 50 μm sections and immediately placed in 4% PFA for fixation overnight for large-scale 3D imaging. The frozen tissue left over, if any, after sectioning was used for RNA extraction and transcriptomic analysis described below. Therefore, all tissues used in transcriptomics had also tissue cytometry performed, except for patient **17**, where the tissue was too small to be sectioned for imaging and was used entirely for RNA analysis.

### Immunofluorescence staining and large-scale 3D confocal imaging

For immunofluorescence analysis, 50 μm sections were washed in PBS and then blocked in 10% normal donkey serum for 2 h at RT, followed by primary antibody incubation at RT overnight and permeabilization with Triton X at 0.2% ^46, 47^. We used the following antibodies for detecting the target proteins: Aquaporin 1 (AQP1, Santa Cruz Biotechnology; sc-9878), myeloperoxidase (MPO, Abcam; ab5690), CD68 (Agilent; M087601), Citrulline H3 (OriGene Technologies Inc; AM10179PU-N). After washing with PBS, the sections were probed with Alexa dye-conjugated secondary antibodies: donkey anti-goat-488 or anti-mouse-568 and anti-rabbit-647. DAPI was used for staining the nuclei. Subsequently, sections were washed for 5 times 1 h each and then fixed in 4% PFA for additional 15 min. After a final wash for 30 mins, sections were mounted on a glass slide using fluoromount medium (Sigma Aldrich; F-4680). Images were sequentially acquired in four separate channels using Leica SP8 confocal microscope as whole volume stacks using 20X NA 0.75 objective with 1.0-*μ*m spacing. Stacks were stitched using Leica LAS X software to generate large scale 3D images. 3D image rendering was performed using Voxx V2.09 software (http://www.indiana.edu/~voxx/). A negative control without primary antibody was used to ensure the absence of non-specific binding of secondary antibodies. Microscope settings were identical between imaging sessions of each specimen.

### 3D tissue cytometry

3D tissue cytometry was performed on image volumes using volumetric tissue exploration and analysis (VTEA) software which allows volumetric cell identification on the basis of 3D nuclear segmentation ^7^. Segmentation settings were adjusted to yield the best quality, which was visually verified by sampling random fields within each image stack. Fluorescence from MPO, CD68 or H3-citrulline associated with nuclei by a 3D morphological grow routine were displayed on a scatterplot as individual points. This method allows gating of specific cell populations based on fluorescence intensities, and mapping of the gated cells directly on the image with the corresponding statistics (Supplemental Fig. S1). Direct visualization of the gated cells on the image also allows validation of the gates, which was performed individually on random fields for each large-scale image volume. For spatial distribution analysis, regions of interest (ROIs) were manually drawn over collecting duct identified by morphological features on the stained tissue. Distance maps corresponding to the ROIs were built using ImageJ Euclidian distance map API, and neutrophil position coordinates were correlated to corresponding locations on the distance map. Randomly generated position coordinates using a random number generator function (Gaussian) were used for comparison. A two sample t-test was used to compare the histogram distributions.

### RNA extraction, library preparation and sequencing

Total RNA extraction was performed using RNeasy Plus Micro Kit (Qiagen). Remaining OCT frozen biopsies were dissolved in RLT buffer (Qiagen) and lysed. Equal volume of 75% ethanol was added, and the rest of the protocol was followed according to the manufacturer’s instructions. Extracted RNA was dissolved in RNase free water. The concentration and quality of total RNA samples were first assessed using Agilent 2100 Bioanalyzer. An RIN (RNA Integrity Number) of 5 or higher was required to pass the quality control. 1 nanogram of RNA per sample was used to prepare dual-indexed strand-specific cDNA libraries using Clontech SMARTer RNA Pico Kit v2. The resulting libraries were assessed for quantity and size distribution using Qubit and Agilent 2100 Bioanalyzer. Two hundred picomolar pooled libraries were utilized per flowcell for clustering amplification on cBot using HiSeq 3000/4000 PE Cluster Kit and sequenced with 2×75bp paired-end configuration on HiSeq4000 (Illumina) using HiSeq 3000/4000 PE SBS Kit. A Phred quality score (Q score) was used to measure the quality of sequencing. More than 90% of the sequencing reads reached Q30 (99.9% base call accuracy).

### Sequence alignment and gene counts

The sequencing data were first assessed using FastQC (Babraham Bioinformatics, Cambridge, UK) for quality control. Then all sequenced libraries were mapped to the human genome (UCSC mm10) using STAR RNA-seq aligner ^48^ with the following parameter: “--outSAMmapqUnique 60”. The reads distribution across the genome was assessed using bamutils (from ngsutils) ^49^. Uniquely mapped sequencing reads were assigned to hg38 refGene genes using featureCounts (from subread) ^50^ with the following parameters: “-s 2 ‒p ‒Q 10”. Quality control of sequencing and mapping results was summarized using MultiQC ^51^. Genes with read count per million (CPM) > 0.5 in more than 4 of the samples were kept. The data were normalized using TMM (trimmed mean of M values) method. Differential expression analysis was performed using edgeR ^52, 53^. False discovery rate (FDR) was computed from p-values using the Benjamini-Hochberg procedure. Differentially expressed genes (uncorrected p<0.01) were subjected to Ingenuity Pathway Analysis (IPA) to determine significant gene clustering (p<0.05) within signaling pathways or other disease annotations (Ingenuity Systems, http://www.ingneuity.com). Volcano plots were generated using the Shiny Transcriptome Analysis Resource Tool (START) ^54^. Heat maps were generated using R software (version 3.4.2)

### Proteomic studies

Kidney stone samples were processed as two technical replicates when available. A total of 10 brushite samples (5 patients) and 9 CaOx samples (5 patients) were analyzed. Kidney stone samples, each >100 mg dry weight, were pulverized and proteins were isolated as previously described and analyzed using a label-free LC-MS/MS method ^6^. Briefly, stones were powdered to the consistency of fine flour. 100 mg of stone powder were combined with 1.0 ml of freshly prepared 8 M urea and 10 mM DTT. Mixture was sonicated and tubes were incubated overnight at RT on orbital shaker (200rpm). After centrifugation, the supernatant was collected and the protein concentration was determined by Bradford assay. A 100 μg aliquot of each sample was concentrated and volume was adjusted to 200 μl (4M urea) and then reduced and alkylated by triethylphosphine and iodoethanol. Reduced/alkylated sample was dried by SpeedVac and reconstituted in 100 μl of NH_4_HCO_3_. 20μg/ml trypsin solution was added into 150 μl aliquot for overnight digestion. Digested samples were inspected using a ThermoScientific Orbitrap Velos Pro hybrid ion trap-Orbitrap mass spectrometer coupled with a Surveyor auto sampler and MS HPLC system (ThermoScientific). The data were collected in the “Data dependent MS/MS” mode of Fourier transform-ion trap (MS-MS/MS) with the electrospray ionization interface using normalized collision energy of 35%. The acquired data were searched against the UniProt protein sequence database of HUMAN using SEQUEST algorithms in Bioworks. Identified proteins were validated by PeptideProphet and ProteinProphet using TransProteomic Pipeline. After validation protein probability of ≥ 0.9000 and peptide probability of ≥ 0.8000 were reported. Protein quantification was determined using a label free quantification software package, IdentiQuantXL™ ^6^. Data were analyzed using the statistical package in MetaboAnalyst 4.0 (https://www.metaboanalyst.ca/MetaboAnalyst/faces/home.xhtml). Heat maps were generated using log-transformed normalized data. Additionally, uncorrected p-values <0.05 were applied to filter affected genes/proteins for function, pathway and network analysis using IPA to determine significant enrichment of canonical pathways and gene clustering (p<0.05) between the two groups.

### Statistical Analysis

The results are presented as the means ± SD. Unless specified otherwise, the significant difference between the experimental groups was assessed by two-tailed *t*-test. For the cytometry results, statistical analyses and graphing were performed using GraphPad Prism (La Jolla, CA), considering p values < 0.05 as statistically significant.

## Author contributions

Experiment design: MSM, SW, FW, JCW, TMEA

Surgical studies: JL, AK, SB, JCW

Clinical studies: EMW, FC, JL, AK, KB, SB, JCW, TMEA

Data collection: MSM, SW, JL, AK, FW, SB, KB, JCW, TMEA

Data analysis: MSM, SW, SK, FW, EMW, FC, APE, KB, DB, JCW, TMEA

Manuscript preparation and editing: all authors

## Acknowledgements

This work was supported by grants from the National Institute of Diabetes and Digestive and Kidney Diseases (NIDDK): P01DK056788 and P30DK079312, and National Institute of Health-Director’s Office: S10OD016208.

## Supplemental data

**Supplemental Figure S1.**
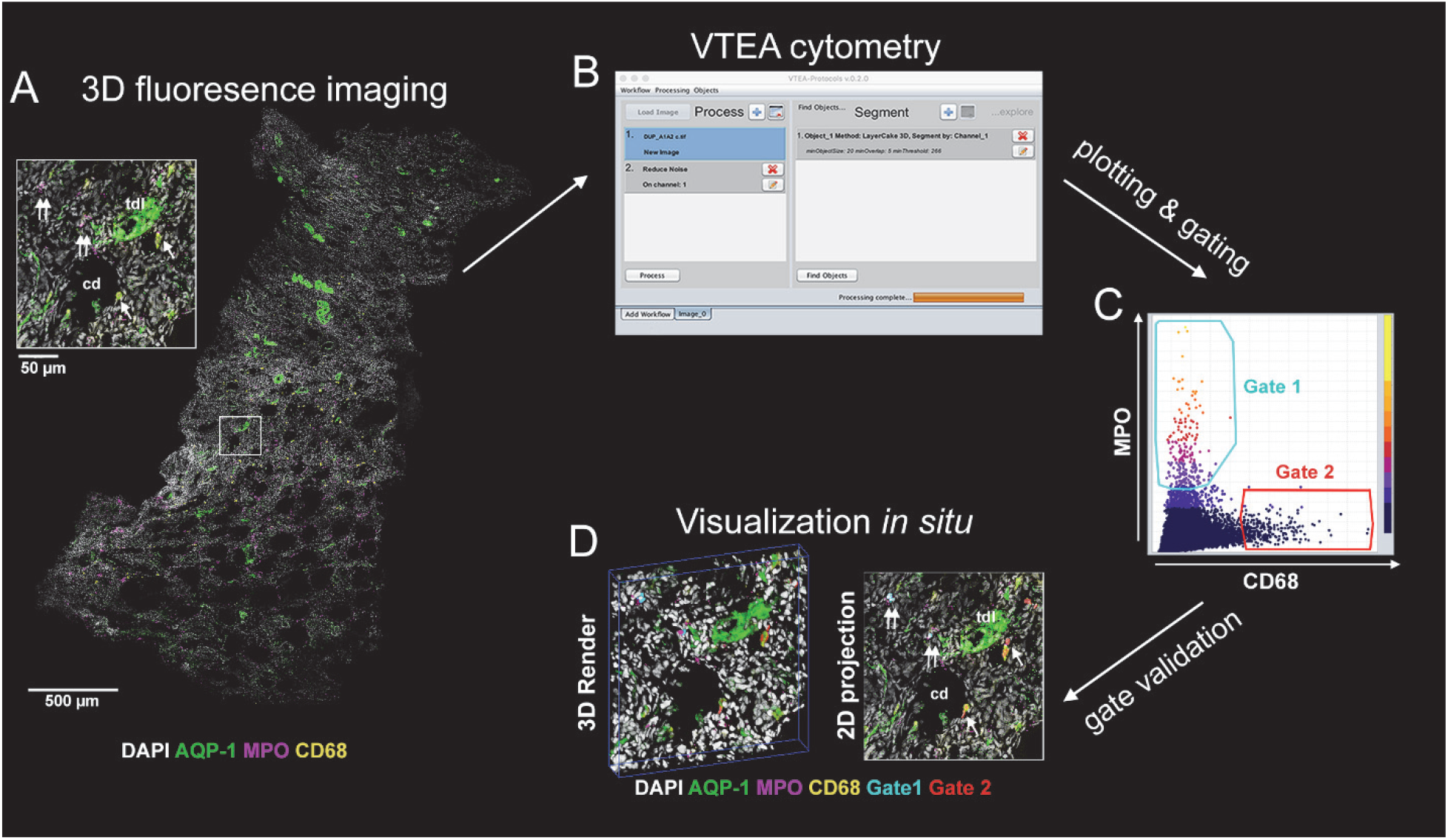
Volumetric Tissue Exploration and Analysis (VTEA) tool for 3D cytometry: (A) 50-μm thick papilla section was imaged using confocal microscope. (B) Four channel-imaged volume was loaded into Image/FIJI. 3D nuclei segmentation was performed by VTEA and (C) VTEA generates scattered plot where each dot represents measured parameters for a single cell. Cytometric events were gated using rectangle or free hand tool which was able to identify subpopulation of cells. Gated cells were highlighted as nuclear overlays in the image volume to identify their spatial distribution (D). This allows immediate validation of the gates used on the scatter plots.

## Legends for supplemental files

**Supplemental data file 1:** Summary file for transcriptomics with differential expression analysis and summary statistics

**Supplemental data file 2:** Summary file for raw data from label free quantitative proteomics for proteins identified in each sample

**Supplemental data file 3:** Neutrophil proteins identified in this study and 5 other publications

